# Autophagy Suppresses CCL2 to Preserve Appetite and Prevent Lethal Cachexia

**DOI:** 10.1101/2025.02.20.638910

**Authors:** Maria Ibrahim, Maria Gomez-Jenkins, Adina Scheinfeld, Zhengqiao Zhao, Eduardo Cararo Lopes, Akshada Sawant, Zhixian Hu, Aditya Dharani, Michael Sun, Sarah Siddiqui, Emily T. Mirek, Johan Abram-Saliba, Edmund C. Lattime, Xiaoyang Su, Tobias Janowitz, Marcus D. Goncalves, Steven M. Dunn, Yuri Pritykin, Tracy G. Anthony, Joshua D. Rabinowitz, Eileen White

## Abstract

Macroautophagy (autophagy hereafter) captures intracellular components and delivers them to lysosomes for degradation and recycling^1^. In adult mice, autophagy sustains metabolism to prevent wasting by cachexia and to survive fasting, and also suppresses inflammation, liver steatosis, neurodegeneration, and lethality^2,3^. Defects in autophagy contribute to metabolic, inflammatory and degenerative diseases, however, the specific mechanisms involved were unclear ^4^. Here we profiled metabolism and inflammation in adult mice with conditional, whole-body deficiency in an essential autophagy gene and found that autophagy deficiency altered fuel usage, and reduced ambulatory activity, energy expenditure, and food intake, and elevated circulating GDF15, CXCL10, and CCL2. While deletion of *Gdf15* or *Cxcl10* provided no or mild benefit, deletion of *Ccl2* restored food intake, suppressed cachexia and rescued lethality of autophagy-deficient mice. To test if appetite suppression by CCL2 was responsible for lethal cachexia we performed single nucleus RNA sequencing of the hypothalamus, the center of appetite control in the brain. Notably, we found that autophagy deficiency was specifically toxic to PMCH and HCRT neurons that produce orexigenic neuropeptides that promote food intake, which was rescued by deficiency in CCL2. Finally, the restoration of food intake via leptin deficiency prevented lethal cachexia in autophagy-deficient mice. Our findings demonstrate a novel mechanism where autophagy prevents induction of a cachexia factor, CCL2, which damages neurons that maintain appetite, the destruction of which may be central to degenerative wasting conditions.

**Key points of paper:**

1) Autophagy-deficient mice have reduced food intake, systemic inflammation, and cachexia
2) CCL2, but not GDF15 or CXCL10, induces lethal cachexia caused by autophagy defect
3) Autophagy-deficient mice have CCL2-dependent destruction of appetite-promoting neurons in the hypothalamus
4) Leptin deficiency restores appetite and rescues lethal cachexia in autophagy-deficient mice
5) Autophagy-deficient mice die from cachexia mediated by appetite loss
6) Degenerative conditions due to impaired autophagy are caused by the inflammatory response to the damage
7) Targeting CCL2 may be a viable approach to prevent degenerative wasting disorders

## Main

Autophagy is regulated by the autophagy-related genes (*Atg*) that function to assemble autophagosomes and capture cargo including intracellular proteins and organelles, for degradation and recycling. Macromolecules produced by autophagy recycling support metabolism and eliminate damaged proteins and organelles thereby suppressing inflammation^5,6^. Autophagy recycling is essential for cell survival in mammals during the absence of nutrients. *Atg5*- or *Atg7*-deficient mice are born developmentally normal but fail to survive the neonatal starvation period due, in part, to nutrient insufficiency^1,7^. Moreover, fasting is lethal to adult mice with conditional, whole-body deletion of *Atg5* or *Atg7* due to hypoglycemia and wasting of muscle and adipose tissue characteristic of cachexia^2,3^. Thus, the evolutionary conserved function of autophagy is sustaining metabolic homeostasis and survival to nutrient deprivation.

Autophagy is also important in the fed state, however, the specific mechanisms are unclear. Conditional ablation of *Atg5* or *Atg7* in adult mice leads to liver inflammation and neurodegeneration, and also weight loss, adipose tissue lipolysis and muscle atrophy, and body wasting characteristic of cachexia^2,3,8^. The lifespan of adult mice with conditional deficiency in autophagy is limited to less than 3 months^2,3^. Interestingly, neuronal-specific *Atg5* expression rescues the neonatal lethality of *Atg5*-deficient mice^7^, however, the aspect of neuronal function that is required to enable survival and if or how this is related to cachexia is unknown^9^. We found that autophagy suppresses CCL2 thereby preserving hypothalamic neurons and food intake, which prevents lethal cachexia. Thus, CCL2 is a cachexia factor responsible for hypothalamic neuron degeneration leading to anorexia and death.

## Results

### Autophagy deficiency leads to weight loss, altered body composition, liver inflammation, and cachexia

To investigate the role of autophagy in whole-body metabolism and cachexia, we analyzed body composition in conditional whole-body *Atg7*-deficient (*Atg7^Δ/Δ^*) compared to autophagy-intact (*Atg7^+/+^)* mice. At ten weeks post-deletion, *Atg7^Δ/Δ^* mice showed consistent reduction in body weight and a greater percent decrease from their initial body weight (Fig. 1a, S1a). *Atg7^Δ/Δ^* mice also exhibited reduced lean mass, progressive fat mass depletion (Fig. 1b-c) with lower weights of white and brown adipose tissue, as well as soleus and gastrocnemius plantaris muscles, compared to *Atg7^+/+^* mice (S1b-c). Previous studies have shown that short-term conditional *Atg7* deletion results in liver inflammation, steatosis, and hepatomegaly^2,10,11^. At ten weeks post-deletion, when liver weight was excluded from body weight, a further reduction in weight was observed in *Atg7^Δ/Δ^* mice (Fig. 1d). Thus, the reduction in body weight seen in *Atg7^Δ/Δ^* mice is an underestimate of body wasting due to the enlarged liver. In *Atg7^Δ/Δ^* mice, serum levels of liver enzymes alanine aminotransferase (ALT) and aspartate aminotransferase (AST) were elevated, indicating liver inflammation and impaired function (Fig. S1d). To explore the relationship between weight loss, systemic inflammation, and autophagy deficiency, we performed RNA sequencing on livers from *Atg7^Δ/Δ^* mice and compared them to *Atg7^+/+^* controls (Fig. S1e). GSTA1, GSTM1, and GSTM3 are genes encoding Glutathione S-Transferases (GSTs), which play a key role in oxidative stress response and detoxification by conjugating reduced glutathione to toxins, were significantly upregulated in the livers of *Atg7^Δ/Δ^*mice. Moreover, expression of two pro-inflammatory cytokines, CXCL10 and CCL2 were upregulated in *Atg7^Δ/Δ^* mice liver consistent with the known role of autophagy in suppressing damage and inflammation in the liver ^2,12^.

**Figure 1:**
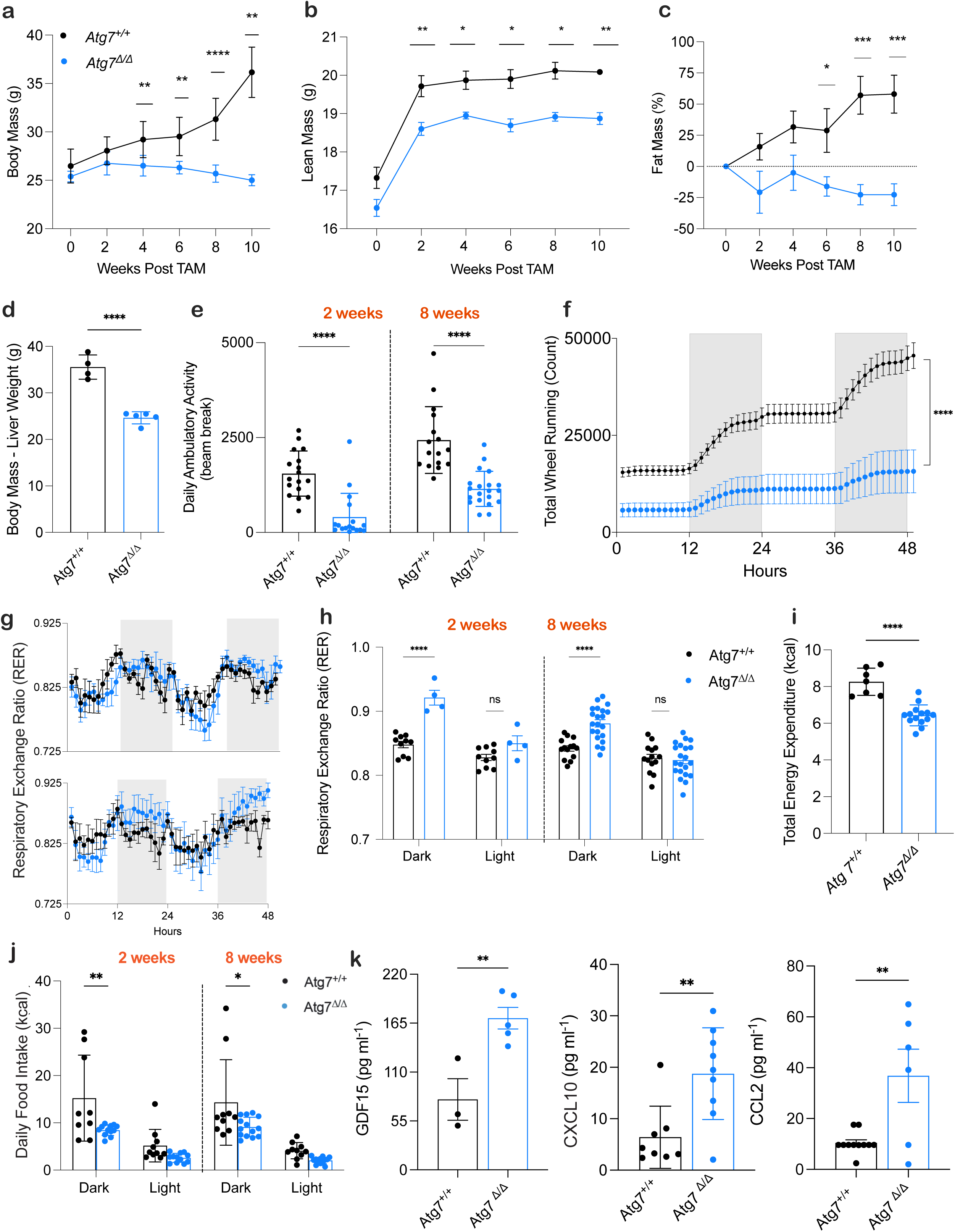
Systemic metabolic impairment due to loss of autophagy causes cachexia. **a,** Mouse body mass post TAM injection in *Atg7^+/+^* mice (*n* = 5) and *Atg7^Δ/Δ^* mice (*n* = 7) **b,c** Lean mass in grams and fat mass percentage loss post TAM injection in *Atg7^+/+^* mice (*n* = 5) and *Atg7^Δ/Δ^*mice (*n* = 5). Body composition was measured by EchoMRI. All data are mean ± s.e.m. \**P* < 0.05, \*\**P* < 0.01, \*\*\**P* < 0.01, \*\*\*\**P* < 0.0001 using a two-sided Student’s *t*-test. **d,** Body mass subtracted by liver weight at 10 weeks post TAM in *Atg7^+/+^* mice (*n* = 4) and *Atg7^Δ/Δ^* mice (*n* = 5) **e**–**j,** Mice were housed in Promethion metabolic cages (*n* = 4–21/group). Shaded regions represent the dark cycle from 19:00 pm to 7:00 am. **e,** daily ambulatory activity at 2- and 8-weeks post TAM. **f,** total wheel running at 2 weeks post TAM. **g,** Hourly mean of RER at 2- and 8-weeks post TAM. **h,** Overall hourly means of RER at 2- and 8-weeks post TAM. **i**, Total energy expenditure. **j,** daily food intake. **k,** Serum (GDF15 ELISA) and cytokine and chemokine profiling (CXCL10 and CCL2) (*n* = 5–11/group) of *Atg7*^+/+^ and *Atg7*^Δ/Δ^ mice.

To monitor metabolic activity in *Atg7^Δ/Δ^* and *Atg7^+/+^*mice they were assessed in metabolic cages every two weeks post-deletion. Conditional whole-body *Atg7*-deficient mice and mice lacking *Atg7* specifically in the central nervous system present with abnormal limb-clasping reflexes and behavioral defects^2,13,14^. Accordingly, we measured behavioral activity through ambulatory activity and total wheel running and found *Atg7^Δ/Δ^* mice displayed lower activity compared to *Atg7^+/+^* mice (Fig. e-f). Ambulatory activity analysis revealed a significant decrease at each timepoint in *Atg7^Δ/Δ^* mice (Fig. S1f). These findings are consistent with neurodegeneration attributed to deficient autophagy^15^.

The respiratory exchange ratio (RER), which represents the ratio of produced CO₂ to consumed O₂ is reflective of the major types of macronutrients being metabolized (lipids vs. carbohydrates). Hourly RER plots at 2- and 8-weeks post-deletion revealed that *Atg7^Δ/Δ^* mice exhibited higher RER values compared to *Atg7^+/+^*mice during the dark phase when mice are active (Fig. 1g). RER analysis across both light and dark phases showed that this increase was statistically significant during the dark phase at each time point (Fig. 1h, S1g). These findings suggest a shift away from fat and/or toward carbohydrate (glucose) utilization as the preferred fuel source. This observation aligns with previous studies showing glycogen depletion in the liver of adult mice with conditional autophagy deficiency^2,16^ and may indicate a compensatory reliance on alternative nutrient sources due to a metabolic deficit. In contrast, no differences were observed between *Atg7^Δ/Δ^* and *Atg7^+/+^*mice during the light phase for RER when mice are inactive (Fig. 1h, S1g).

Involuntary weight loss is often linked to reduced appetite and/or increased energy expenditure, which disrupts energy homeostasis. Total energy expenditure (TEE), which encompasses basal metabolism, thermoregulation, physical activity, and the thermic effect of food intake, was reduced in *Atg7^Δ/Δ^* mice, failing to account for weight loss (Fig. 1i). Remarkably, we found that *Atg7^Δ/Δ^* mice exhibited significantly lower food intake compared to *Atg7^+/+^*mice (Fig. 1j), surprising given that they are intolerant to fasting, but possibly explaining loss of lean and fat mass. As cytokines and chemokines can regulate appetite^17,18^ we measured these factors in the serum (Fig. S1h). We found three factors that were significantly upregulated in the circulation of *Atg7^Δ/Δ^* mice compared to *Atg7^+/+^* mice: Growth differentiation factor 15 (GDF15), C-X-C motif chemokine ligand 10 (CXCL10), and C-C motif ligand 2 (CCL2) (Fig. 1k). Thus, autophagy-deficient mice display a cachexia-like syndrome, including loss of body weight, appetite, and wasting of muscle and fat, and increased levels of circulating cytokines and chemokines.

### CCL2 is the dominant factor responsible for lethal cachexia in autophagy-deficient mice

To determine if GDF15, CXCL10 or CCL2 contribute to cachexia in autophagy-deficient mice, we generated double knockout mice for each factor on the conditional *Atg7^Δ/Δ^* background. GDF15 is a hormone known to reduce food intake ^19,20^ by causing food aversion via signaling through specific neurons in the area postrema and the nucleus of the solitary tract that express its receptor GFRAL^21,22^. To determine whether GDF15 modulates the lethality in autophagy deficiency, we generated mice with constitutive deficiency in *Gdf15*^23,24^ and crossed them to *Ubc-Cre^ERT2/+^; Atg7^flox/flox^* mice to generate *Gdf15^−/−^; Ubc-Cre^ERT2/+^; Atg7^flox/flox^* mice. Tamoxifen (TAM) administration was used to delete *Atg7* in the presence and absence of GDF15 (Fig. S1i). The loss of GDF15, however, neither rescued the lethality caused by autophagy deficiency (Fig. S1i) nor did it impact any other obvious phenotype.

CXCL10 is a chemokine induced in association with metabolic diseases^25^ and infection^25–27^. To investigate whether CXCL10 induction contributed to altered metabolism and reduced survival caused by autophagy deficiency, mice with constitutive deficiency in *Cxcl10*^27^ were crossed with *Ubc-Cre^ERT2/+^; Atg7^flox/flox^* mice to generate *Cxcl10 ^−/−^; Ubc-Cre^ERT2/+^; Atg7^flox/flox^* mice. TAM administration was used to delete *Atg7* in the presence and absence of CXCL10 (Fig. S1j). *Cxcl10^−/−^;Atg7^Δ/Δ^* mice demonstrated a small but significant improvement in survival compared to *Atg7^Δ/Δ^* mice, with median survival increased from 64 days to 134 days (Fig. S1j). These results suggest that CXCL10 modestly extends survival, but neither GDF15 nor CXCL10 deficiency is sufficient to substantially rescue weight loss, food intake, and lethality resulting from autophagy deficiency.

CCL2 is a chemokine that recruits monocytes, macrophages, and other immune cells to sites of injury or infection^23,28^. It has been previously implicated in cancer-induced cachexia^29^, specifically metabolic changes in muscle and white adipose tissue (WAT)^30^. To test whether CCL2 impacts the survival of mice lacking autophagy, mice with constitutive deficiency in *Ccl2*^23^ were crossed with *Ubc-Cre^ERT2/+^; Atg7^flox/flox^* mice to generate *Ccl2^−/−^; Ubc-Cre^ERT2/+^; Atg7^flox/flox^* mice. TAM administration was used to delete *Atg7* in the presence and absence of CCL2 (Fig. 2Sa). While *Atg7^Δ/Δ^* mice survived less than three months^2^, the loss of CCL2 completely rescued lethality induced by autophagy deficiency (Fig. 2a). Notably, loss of CCL2 did not induce major alterations in the cytokine and chemokine profile comparing *Atg7^Δ/Δ^* and *Atg7*^+/+^ mice, suggesting that it may function directly (Fig. S2b).

**Figure 2:**
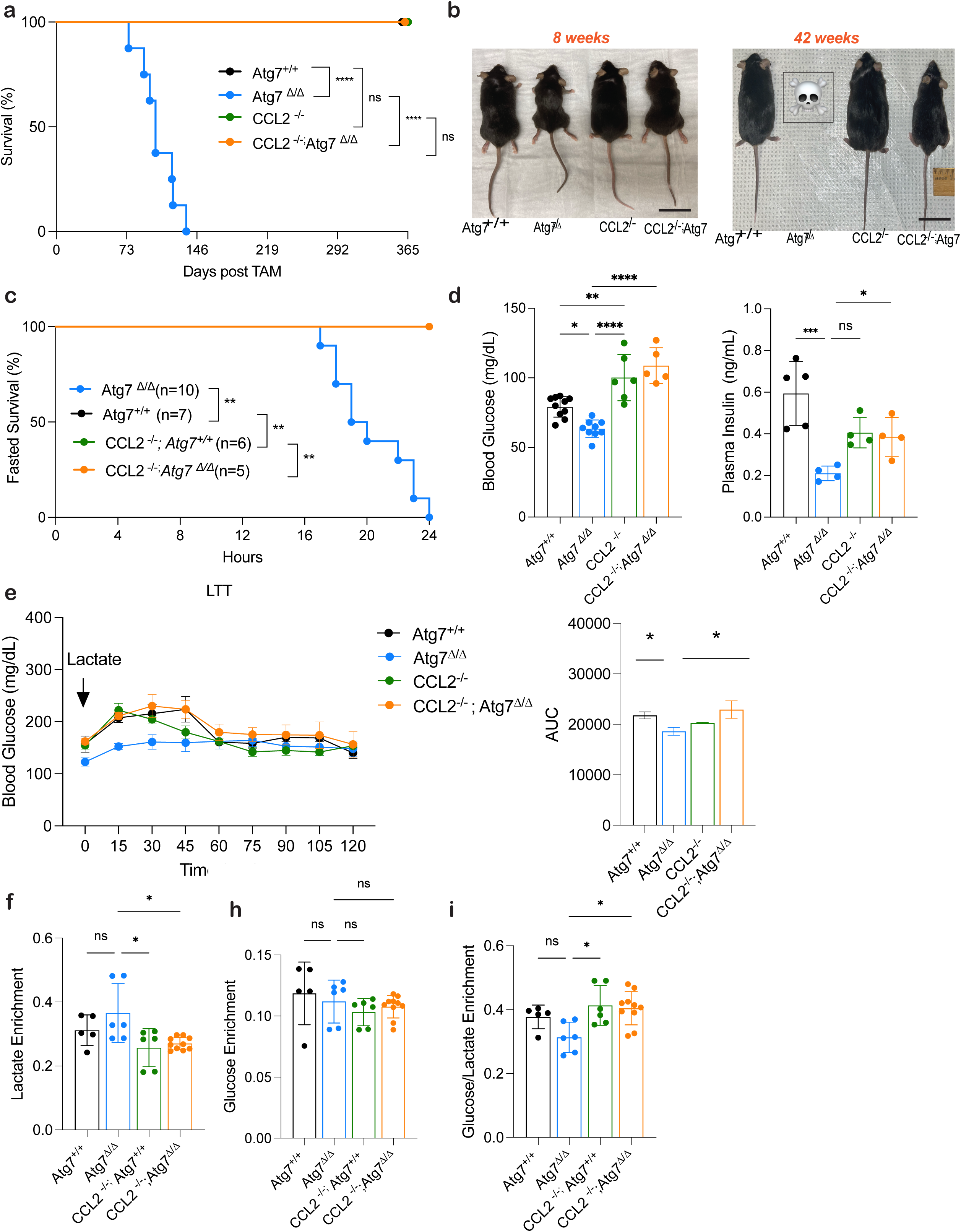
Induction of CCL2 contributes to lethality during autophagy deficiency. **a,** Kaplan-Meier survival curve of *Atg7^+/+^*, *Atg7^Δ/Δ^*, *Ccl2^-/-^*, and *Ccl2^−/−^;Atg7^Δ/Δ^* mice. **b,** Representative images of *Atg7^+/+^*, *Atg7^Δ/Δ^*, *Ccl2^-/-^*, and *Ccl2^−/−^;Atg7^Δ/Δ^* mice at 8- and 42-weeks post TAM injection. **c,** Kaplan-Meier 24 hours fasting survival curve of *Atg7^+/+^*, *Atg7^Δ/Δ^*, *Ccl2^-/-^*, and *Ccl2^−/−^;Atg7^Δ/Δ^* mice 10 days post–TAM. **d,** Blood glucose and plasma insulin measurements collected at 16-hour post fast. **e,** Blood glucose following an intraperitoneal lactate tolerance test. Area under curve calculated from individual blood glucose traces. (*) *P* < 0.05; (***) *P* < 0.001; (****) *P* < 0.0001; (n.s.) not significant (unpaired *t*-test). **f-i,** Statistical analysis of the main altered metabolites enrichment in plasma of *Atg7^+/+^*, *Atg7^Δ/Δ^*, *Ccl2^-/-^*, and *Ccl2^−/−^;Atg7^Δ/Δ^* mice after *in vivo* ^13^C lactate tracing at 2 weeks post deletion. **f,** Lactate enrichment **h,** glucose enrichment **i,** ratio glucose/lactate enrichment. For all graphs the *P* values were determined using one-way ANOVA. *P* values are indicated as ≤0.05*, ≤0.01**, ≤0.001***, and ≤0.0001****.

To determine how eliminating CCL2 rescued lethality of autophagy-deficient mice we characterized the phenotypes of the four mouse genotypes. Histologic examination of tissues by H&E showed that CCL2 deficiency mitigated tissue damage resulting from loss of autophagy including liver inflammation, depletion of the lipid content of white adipose tissue (WAT) and brown adipose tissue (BAT), and atrophy of skeletal muscle (Fig. S2c). Notably, the physical appearance of cachexia, the increased livers weights, and the impaired liver function was diminished in the *Ccl2^−/−^;Atg7^Δ/Δ^* compared to *Atg7^Δ/Δ^* mice (Fig. 2b, Sd-e). While *Atg7^Δ/Δ^* mice develop evidence of severe hepatic dysfunction, as assessed by hyperbilirubinemia, low triglyceride levels, and low blood urea nitrogen (BUN), the *Ccl2^−/−^; Atg7^Δ/Δ^* mice were protected (Fig. S2f). Together, these results suggested that CCL2 plays a crucial role in maintaining survival and preventing tissue damage upon loss of autophagy.

CCL2 induction is associated with weight loss including depleting muscle and adipose tissue while also inducing liver steatosis^29^, particularly during systemic inflammation^31^ and neurodegeneration^32^ similar to what we observed in autophagy-deficient animals. As such, therapeutic targeting of CCL2 with antibodies was attempted, but unfortunately without success^33^. To test if inhibiting CCL2 with an antibody was equivalent to genetic *Ccl2* deficiency we regenerated the C1142 monoclonal antibody (mAb) proposed to neutralize circulating CCL2^34,35^. Following TAM-induced autophagy deficiency, mice were treated with either C1142 mAb or an IgG control antibody. While *Atg7^Δ/Δ^* mice treated with either C1142 mAb or IgG mAb showed no difference in survival (Fig. S2g), the *Atg7^Δ/Δ^* mice treated with C1142 mAb showed partial rescue of body weight over time compared to IgG mAb (Fig. S2h). However, this result was due to an increase in lean mass from further increased hepatomegaly in *Atg7^Δ/Δ^* mice rather than a prevention of adipose and skeletal muscle wasting (Fig. S2i). These data suggest that an antibody directed against a CCL2 peptide does not phenocopy genetic deletion of *Ccl2*. Notably, CCL2 levels in the liver of *Atg7^Δ/Δ^* mice treated with C1142 mAb showed a decreasing trend compared to IgG-treated mice, although no significant (Fig. S2j). These observations are in agreement with previous clinical trial observations with therapeutic anti-CCL2 candidates that similarly failed to deplete the chemokine^36^. Moreover, they suggest that previous attempts to target CCL2 with an antibody in vivo were likely ineffective and perhaps counterproductive.

### Loss of CCL2 rescues fasting lethality by preserving liver gluconeogenesis

The loss of CCL2 extends lifespan and attenuates tissue damage in *Atg7^Δ/Δ^* mice (Fig. 2a,S2c). Therefore, we sought to investigate if elevated CCL2 levels also contributed to the fasting-induced mortality due to hypoglycemia in autophagy-deficient mice^2,3^ Mice were subjected to fasting (free access to water without food for 24 hours). In contrast to the *Atg7^Δ/Δ^*mice that die upon fasting, *Ccl2^−/−^;Atg7^Δ/Δ^* mice survive (Fig. 2c). Blood glucose and serum insulin levels during fasting were maintained in *Ccl2^−/−^;Atg7^Δ/Δ^*compared to *Atg7^Δ/Δ^* mice, which present with hypoglycemia and reduced insulin levels (Fig. 2d). We hypothesized that elevated blood glucose levels in *Ccl2^−/−^;Atg7^Δ/Δ^* compared to *Atg7^Δ/Δ^*mice resulted from preservation of liver function and the ability to perform gluconeogenesis during fasting. To test this hypothesis, we measured gluconeogenesis by injecting mice with L-Lactate and then measuring the resulting glucose levels in the blood. *Atg7^Δ/Δ^* mice showed impaired ability to utilize lactate for glucose synthesis compared to *Ccl2^−/−^;Atg7^Δ/Δ^* mice, which maintained this capacity (Fig. 2e). To confirm that loss of CCL2 restored hepatic gluconeogenesis in autophagy-deficient mice, we performed *in vivo* ^13^C lactate tracing. The labeled lactate in the plasma of *Atg7^Δ/Δ^* mice was significantly higher as compared to *Ccl2^−/−^;Atg7^Δ/Δ^* mice (Fig. 2f), while the plasma glucose enrichment levels remained unchanged between the groups (Fig. 2h). However, the ratio of glucose to lactate showed that significantly less lactate was being converted to glucose in *Atg7^Δ/Δ^* compared to *Ccl2^−/−^;Atg7^Δ/Δ^* mice (Fig. 2i). These findings demonstrated that *Atg7^Δ/Δ^* mice are unable to efficiently utilize circulating lactate for gluconeogenesis, resulting in reduced blood glucose levels and lethality upon fasting due to hepatic dysfunction. In contrast, *Ccl2^−/−^;Atg7^Δ/Δ^* mice effectively convert lactate to glucose via gluconeogenesis, maintaining blood glucose levels and animal survival during fasting.

### CCL2 deficiency rescues weight and food intake but not fuel utilization or ambulatory activity

In contrast to *Atg7^Δ/Δ^* mice, *Ccl2^−/−^;Atg7^Δ/Δ^* mice maintained body weight, lean mass, and fat mass (Fig. 3a-c) in addition to survival. Interestingly, *Ccl2^-/-^*mice presented with a larger initial body weight, gained significantly more weight compared to the other genotypes, and accumulate larger lipid deposits in adipose tissues compared to *Atg7^+/+^* mice (Fig. 3a-c, S2c). Together these results suggest a role for CCL2 in regulating body composition.

**Figure 3:**
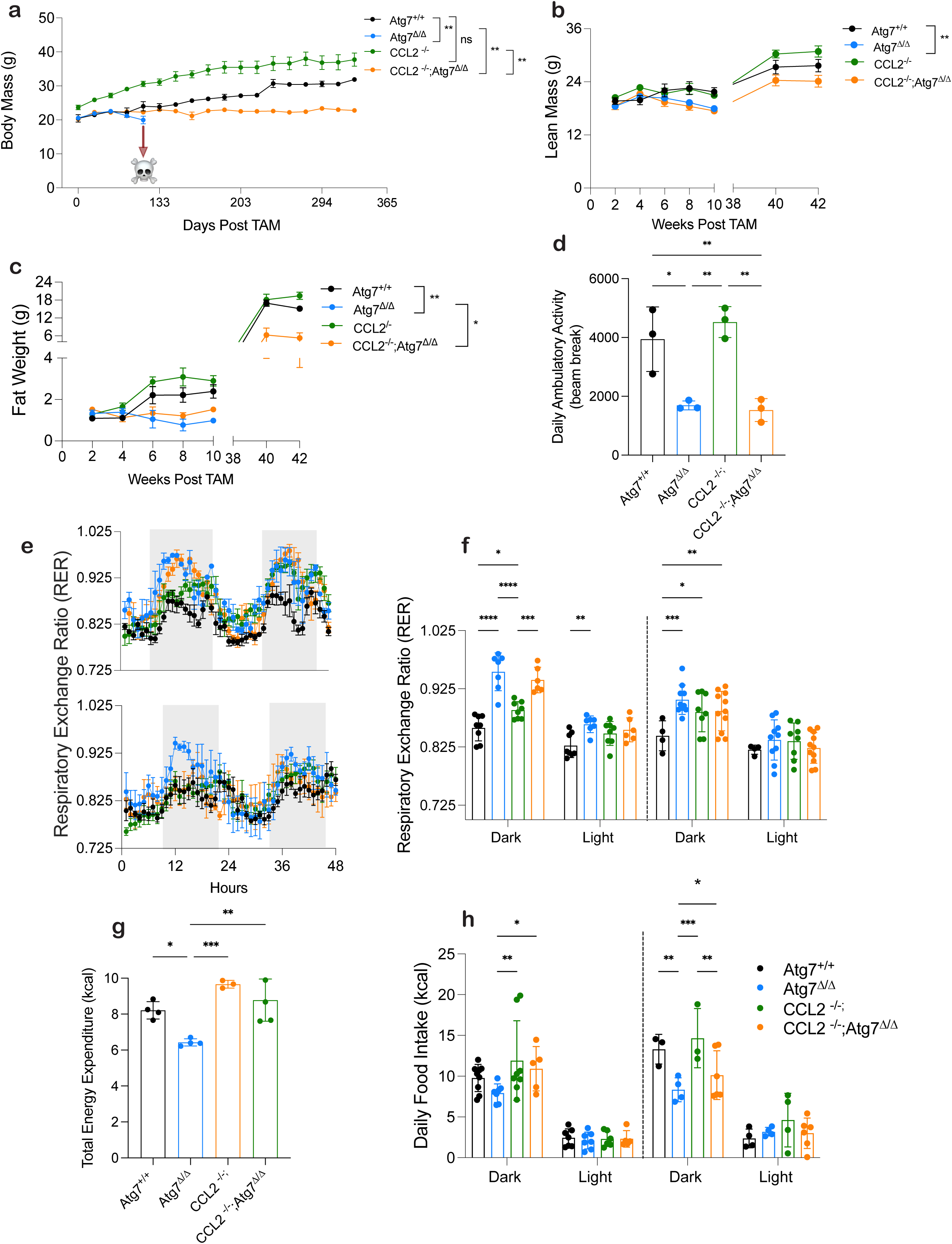
Metabolic Phenotyping shows Loss of CCL2 impacts body composition and food intake. **a,** Mouse body weight post TAM injection in *Atg7^+/+^*, *Atg7^Δ/Δ^* mice, *Ccl2^-/-^* mice, and *Ccl2^−/−^;Atg7^Δ/Δ^* mice over 365 days. **b,c** Lean mass and fat mass over 42 weeks post TAM injection in *Atg7^+/+^*, *Atg7^Δ/Δ^* mice, *Ccl2^-/-^* mice, and *Ccl2^−/−^;Atg7^Δ/Δ^*. Body composition was measured by EchoMRI. All data are mean ± s.e.m. \**P* < 0.05, \*\**P* < 0.01, \*\*\**P* < 0.01, \*\*\*\**P* < 0.0001 using a two-sided Student’s *t*-test. **d–h,** Mice were housed in Promethion metabolic cages (*n* = 4–11/group). Shaded regions represent the dark cycle from 19:00 pm to 7:00 am. **d,** daily ambulatory activity at 2 weeks post TAM. **e,** Hourly mean of RER at 2- and 8-weeks post TAM. **f,** Overall hourly means of RER at 2- and 8-weeks post TAM. **g**, Total energy expenditure. **h,** daily food intake.

Metabolic phenotyping found *Ccl2^−/−^*;*Atg7^Δ/Δ^*and *Atg7^Δ/Δ^* mice showed no significant difference in the RER, suggesting nutrient utilization and preference was similar (Fig. 3e-f). Ambulatory activity also showed no significant difference between *Ccl2^−/−^Atg7^Δ/Δ^* compared to *Atg7^Δ/Δ^*mice (Fig. 3d) and there was partial rescue in progressive motor, ataxia, and behavioral deficits in *Ccl2^−/−^;Atg7^Δ/Δ^* compared to *Atg7^Δ/Δ^* mice (Fig. S2k) and Supplementary Movie S1. Brain histological analyses showed the numbers of pyramidal neurons and Purkinje cells, related to motor function and coordinated movement, were significantly increased in *Ccl2^−/−^;Atg7^Δ/Δ^* compared to *Atg7^Δ/Δ^* mice (Fig. S2l). These results indicated that induction of CCL2 in autophagy deficient mice was not responsible for alternated RER or defective ambulatory activity although there was some preservation of Purkinje cells in the cerebellum and some mitigation of defective hindlimb clasping. Eliminating CCL2, therefore, does not rescue all autophagy-defect related phenotypes and would not be expected to correct cell damage induced by failure of protein and organelle clearance critical to the function of post-mitotic and motor neurons^37^.

As shown above*, Atg7^Δ/Δ^* mice have decreased food intake, TEE, and high levels of CCL2. We therefore measured TEE and food intake in *Ccl2^−/−^* and *CCL2^−/−^; Atg7^Δ/Δ^* mice. Loss of CCL2 restored TEE comparable to *Atg7^+/+^* mice (Fig. 3g). Interestingly, *Ccl2^−/−^* mice exhibited increased food consumption during the dark and light cycle when compared to *Atg7^+/+^* mice (Fig. 3h). Surprisingly, loss of CCL2 significant preserved food intake in *Ccl2^−/−^;Atg7^Δ/Δ^* mice compared to *Atg7^Δ/Δ^* mice at both 2- and 8-weeks post deletion. (Fig. 3h). These results suggest that CCL2 induction in autophagy-deficient mice inhibited appetite, decreased food intake, and disrupted energy homeostasis, which would be potentially lethal as they are intolerant to fasting.

### Eliminating CCL2 rescues loss of appetite-promoting hypothalamic neurons

The ability of CCL2 deficiency to preserve food intake in autophagy-deficient mice suggested that CCL2 may be toxic to neurons in the hypothalamus that express its cognate receptor, CCR2, and produce hormones that regulate food intake^38^. To test this hypothesis, single nucleus RNA sequencing (snRNA-seq) was applied to the hypothalamus from wild-type and *Ccl2^−/−^* mice with and without deletion of *Atg7*. The hypothalami were pooled with four samples per genotype due to the low weight of the tissue. This analysis yielded 20,297 high-quality single-nucleus transcriptomes (Fig. 4a, S3a). Using molecular markers of known hypothalamic regions and cell types^39^, we were able to annotate the major hypothalamic cell type populations for each of the four mouse genotypes (Fig. 4b). We identified 52 clusters that were classified into 28 broad cell types, including astrocytes, fibroblast, oligodendrocytes, GABAergic (GABA) and glutamatergic (GLU) neurons (Fig. 4b). UMAP embedding of each model is also shown (Fig. 4c, S3b). UMAP embedding of each model is also shown (Fig. 4c, S3b). Notably, a cell subpopulation forming Cluster 4 did not match to any known cell types from prior studies (Fig. 4a) ^39^. Cells in Cluster 4 had a higher level of mitochondrial gene expression and lower overall snRNA-seq signal than other clusters, suggesting they were more likely apoptotic (Fig. S3c). Cluster 4 cells were predominately from the *Atg7^Δ/Δ^* hypothalamus (Fig. 4c,d), as compared with the remaining clusters which had a relatively even distribution across the four genotypes. Note that CCL2 expression was predominantly in the fibroblast cluster in *Atg7^Δ/Δ^* mice (Fig. 4e). These findings suggest that Cluster 4 may represent cells in the hypothalamus that are negatively impacted by loss of autophagy and that are restored by co-deletion of *Ccl2*.

**Figure 4:**
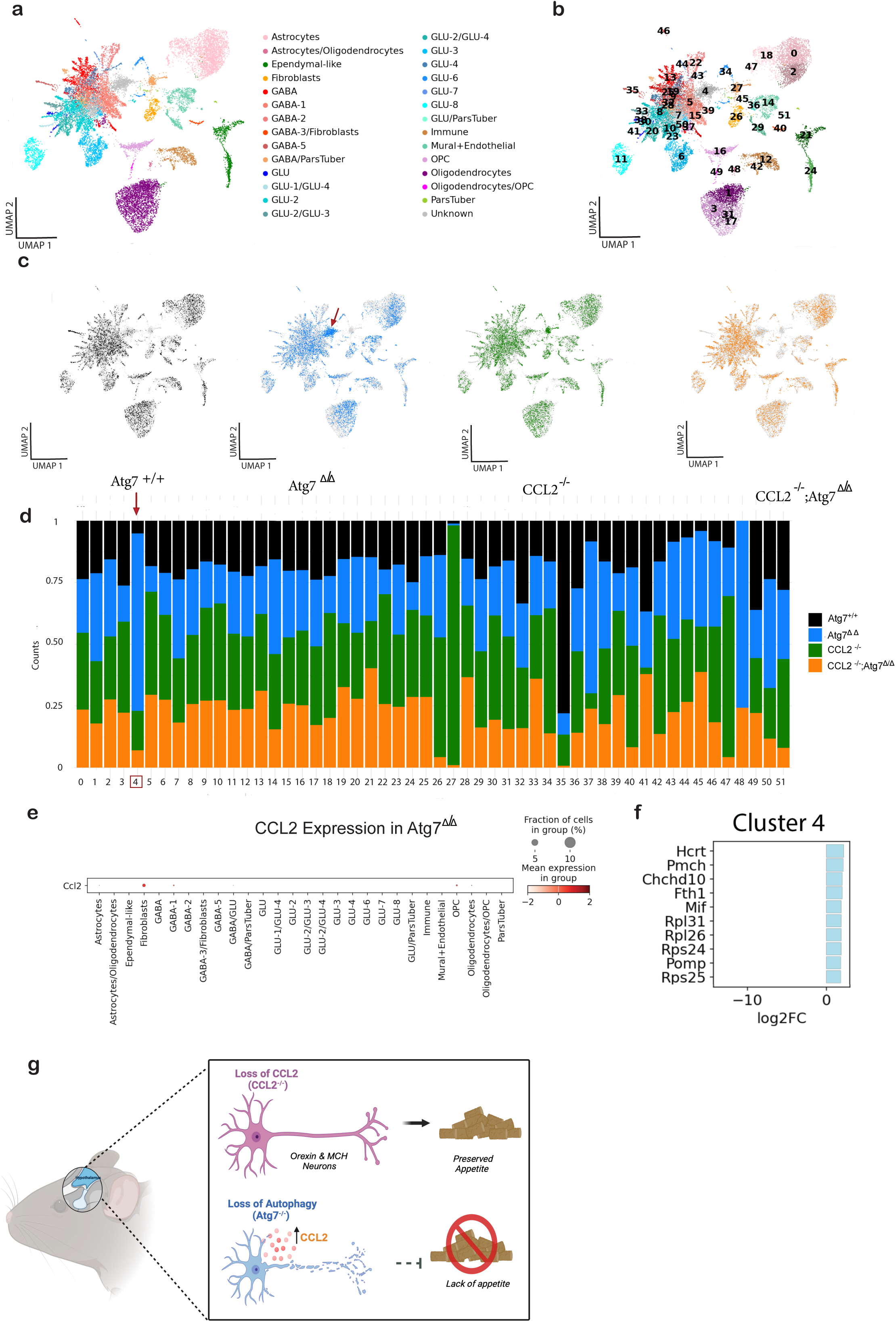
Diversity and proportion of cell types in the scRNA-seq of the hypothalamus from wild-type and *Ccl2^−/−^*mice with and without deletion of *Atg7*. **a,** Uniform Manifold Approximation and Projection (UMAP) of the snRNA-seq data with cell type annotations for *Atg7^Δ/Δ^*, *Ccl2^-/-^*, and *Ccl2^−/−^;Atg7^Δ/Δ^* mice at the 8-wk time point. **b,** UMAP showing 52 clusters that were used to annotate cell types. **c,** UMAP showing the cells separately for *Atg7^Δ/Δ^*, *Ccl2^-/-^*, and *Ccl2^−/−^;Atg7^Δ/Δ^* mice. **d,** Bar plot depicting the cluster composition across the different samples. **e,** Expression of CCL2 across cell types in *Atg7^Δ/Δ^* mice. **f,** Top 10 upregulated genes in Cluster 4. **g,** Schematic of snRNA-seq results due to loss of CCL2.

Positive logFC values in Cluster 4 had significant upregulation of hypocretin (Hcrt), which encodes the neuropeptide orexin, and pro-melanin-concentrating hormone (Pmch), the precursor gene that encodes the neuropeptide melanin-concentrating hormone (MCH) (Fig.4f). Both orexin and pro-MCH are orexigenic hormones that stimulate appetite. To validate the snRNA-seq gene expression analysis from Cluster 4, qRT-PCR analysis was used to measure pro-MCH mRNA expression in the hypothalamus. Hypothalami from *Atg7^Δ/Δ^* mice had decreased mRNA expression, while *Ccl2^−/−^;Atg7^Δ/Δ^* mice had restored mRNA expression similar to *Atg7^+/+^* and *Ccl2^−/−^* mice (Fig. S3d). This data suggested that loss of autophagy leads to CCL2-dependent degradation of cells represented by Cluster 4, which is composed of neurons that produce pro-MCH and orexins, both orexigenic neuropeptides. Thus, the CCL2-induced defective food intake in *Atg7^Δ/Δ^* mice may be due to degradation of neurons that produce positive regulators of appetite, the loss of which may be lethal (Fig. 4g).

### Preservation of appetite rescues survival of autophagy-deficient mice

To test the hypothesis that inhibition of food intake was lethal to autophagy-deficient mice we evaluated if eliminating leptin, an appetite suppressing hormone that signals through the hypothalamus, could rescue their defective food intake and survival. Leptin-deficient humans and mice are obese due to their inability to suppress appetite and food intake^40^. Leptin deficient (*ob/ob)* mice^41^ were crossed with *Ubc-Cre^ERT2/+^;Atg7^flox/flox^* mice to generate *ob/ob;Ubc-Cre^ERT2/+^;Atg7^flox/flox^* mice. TAM administration was used to delete *Atg7* in the presence or absence of leptin. Leptin deficiency rescued lethality of autophagy deficient *ob/ob;Atg7^Δ/Δ^* mice, which survived >250 days post deletion (Fig. 5a). Representative images of each mouse genotype show the weight distribution between respective groups (Fig. 5b). *ob/ob* and *ob/ob;Atg7^Δ/Δ^* mice had similar obese body weights, and fat mass compared to cachectic *Atg7^Δ/Δ^* mice (Fig. 5c,d). Lean mass was comparable between *ob/ob;Atg7^Δ/Δ^* and *Atg7^Δ/Δ^* mice. (Fig. 5e). Additionally, the levels CCL2 were comparable in *ob/ob;Atg7^Δ/Δ^* and *Atg7^Δ/Δ^* mice indicating that leptin deficiency does not rescue survival of autophagy-deficient mice by eliminating CCL2 (Fig. 5f). Leptin deficiency also rescued fasting lethality of autophagy-deficient mice due to the rescue of hypoglycemia and cachexia (Fig. 5g). Lastly, food intake was also rescued in in *ob/ob;Atg7^Δ/Δ^* and *Atg7^Δ/Δ^* mice (Fig. 5h). Thus, autophagy-deficient mice die due to CCL2-mediated suppression of appetite and food intake that can be rescued by increasing appetite and food intake by deleting leptin (Fig. 5i). As autophagy-deficient mice fail to survive fasting, loss of appetite and food intake is lethal.

**Figure 5:**
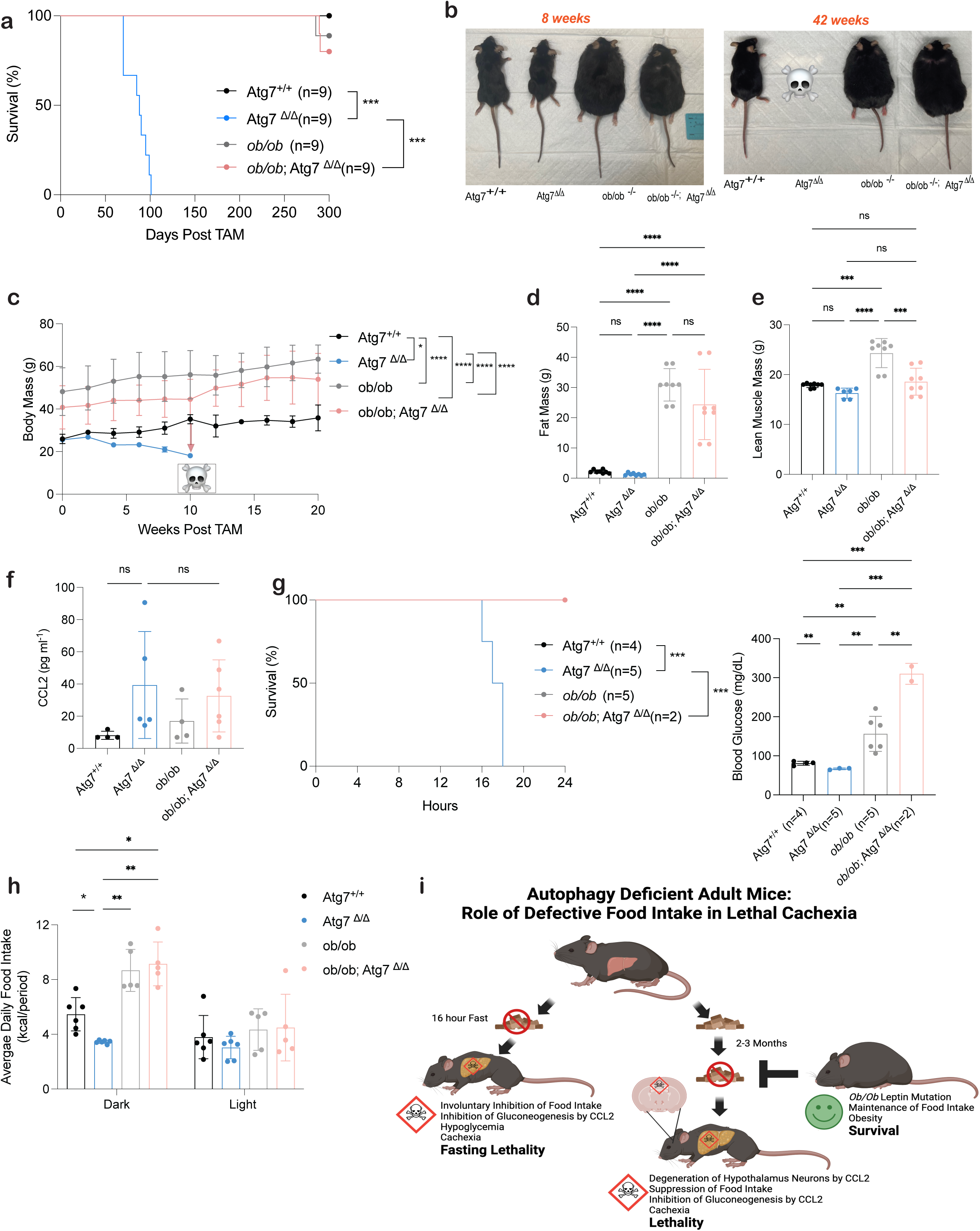
*ob/ob* rescues lethality and weight loss induced by autophagy deficiency. **a,** Kaplan-Meier survival curve of *Atg7^+/+^*, Atg7^Δ/Δ^, *ob/ob*, and *ob/ob*;Atg7^Δ/Δ^ mice. **b,** Representative images of *Atg7^+/+^*, *Atg7^Δ/Δ^* mice, *ob/ob* mice, and *ob/ob*; *Atg7^Δ/Δ^* mice at 8- and 42-weeks post TAM injection. **c,** Mouse body weight post TAM injection in *Atg7^+/+^*, *Atg7^Δ/Δ^* mice, *ob/ob* mice, and *ob/ob*; *Atg7^Δ/Δ^* mice. **d-e,** Fat mass and lean mass loss post TAM injection in mice. Body composition was measured by EchoMRI. All data are mean ± s.e.m. \**P* < 0.05, \*\**P* < 0.01, \*\*\**P* < 0.01, \*\*\*\**P* < 0.0001 using a two-sided Student’s *t*-test. **f,** Serum CCL2 ELISA. **g,** Kaplan-Meier 24 hours fasting survival curve of *Atg7^+/+^*, *Atg7^Δ/Δ^*, *ob/ob*, and *ob/ob*;Atg7^Δ/Δ^ mice 10 days post–TAM. Blood glucose collected at 16-hour post fast. **h,** daily food intake. **k,** Proposed graphical summary of lethality in autophagy deficient mice.

## Discussion

CCL2 is induced in activated microglia in neuroinflammatory diseases and its transgenic expression in mice is sufficient to produce neuronal damage. CCL2 and its receptor CCR2 are associated with STAT2 and IL1β activation and neurodegeneration^32,42,43^, but the mechanisms involved are unclear. CCL2 is also associated with cachexia in cancer models. Administration of CCL2 to mice induces wasting of skeletal muscle^44^ and recruitment of macrophages by CCL2 to tumors promotes cachexia^45^, by unknown mechanisms. The lack of food intake in the *Atg7*-deficient mice is associated with anorexia mediated in the hypothalamus. This is distinct from the effect that GDF15 and the inflammatory cytokine IL-6 that seem to mediate anorexia via receptors in the area postrema.

The chronic loss of the rat MCH-precursor *Pmch* decreases food intake^46,47^ and also affects energy expenditure^46^, thus providing insight into the changed body weight dynamics during chronic loss of *Pmch*. These findings are consistent with loss of autophagy promoting damage, neuroinflammation and CCL2 that destroys MCH-producing orexigenic neurons in the hypothalamus that drive cachexia. Our findings also suggest that targeting CCL2 for degenerative diseases needs to be reexamined due to technical limitations of the approaches in the past. Cachexia is a feature of neurodegenerative and other unresolvable diseases^48,49^. Our findings provide powerful evidence that CCL2 is a cachexic factor that works by suppressing appetite by inhibiting neurons that produce orexigenic peptides. Clear demonstration that CCL2-induced loss of appetite causes lethal cachexia derived from our ability to restore appetite, prevent weight loss and rescue lethal cachexia by eliminating leptin.

Autophagy protects from numerous degradative and inflammatory diseases, and this knowledge has provoked efforts to enhance autophagy for therapeutic benefit^50^. Our findings reveal that much of the damage from autophagy inhibition is surprisingly mediated by CCL2. The orexigenic MCH neurons that are the target of CCL2 express a CCL2 receptor, CCR2^38^, but not CCL2 itself. Thus, loss of autophagy that triggers production of CCL2 occurs in cells other than the HCRT and MCH neurons themselves, perhaps in fibroblasts within the hypothalamus or in activated microglia. These findings also suggest that tissue damage, for example through inhibition of autophagy as shown here, is greatly amplified by the ensuing inflammatory response to that damage. Thus, limiting or resolving the inflammatory response rather than trying to prevent the damage is an alternative approach to mitigate neurodegeneration and other degenerative conditions. Finally, we demonstrate how destructively lethal cachexia can be, as autophagy-deficient mice, despite having several other afflictions, die because they stop eating, illustrating the importance of addressing mechanisms underlying cachexia.

## Author contributions

MI designed, performed genomic data analysis. MGJ, A. Sawant, ECL, and ETM assisted with mice experiments. JA-S and SMD produced and generated C1142 antibody. ZH assisted with maintaining mouse colonies, ear tagging and antibody treatments. AD, MS, and SS assisted with genotyping. JDR, XS, TGA, MDG, and TJ provided data analysis, result interpretation, and contributed valuable suggestions. YP, A. Scheinfeld, and ZZ supported the sequencing, interpreted results, and assisted with writing. EW conceived and supervised the study. All authors read, edited, and approved the manuscript.

## Acknowledgements

This work was supported by the Ludwig Institute for Cancer Research, Ludwig Princeton Branch, Princeton University of EW, and Cancer Grand Challenges CGCSDF-2021\100003 (CRUK) and 1OT2CA278609-01 (NCI) to EW, MDG and TJ, and NIH grant R01 CA163591 to EW, JDR, and NIH grant R01 CA243547 to EW, and NIH grant DP2 AI171161 awarded to Y.P. MI was also supported by postdoctoral fellowship COCR23PRF005 from New Jersey Commission on Cancer Research (NJCCR), MG-J was also supported by 1T32CA257957, ECL was supported by R01 CA243547, and A. Sawant was supported by postdoctoral fellowship DFHS10PPC029 from NJCCR. We thank Rutu Patel, Suman Amjad, Angelina Gruszecki, and Ann Hinrichs for their help with genotyping. SMD and JA-S (LAbCore) were funded directly by Ludwig Cancer Research. We acknowledge support from the Rutgers Cancer Institute Metabolomics Shared Resource (NCI-CCSG P30CA072720-6852), the Biospecimen Repository and Histopathology Service Shared Resource (P30CA072720-5919), and the Immune Monitoring and Flow Cytometry Shared Resource, supported, in part, with funding from NCI-CCSG P30CA072720-6852. The schematic representations were created using BioRender.com. The authors also acknowledge the support from all members of the White laboratory.

## Materials and Methods

### Mouse Models

All animal care was carried out in compliance with Rutgers University Institutional Animal Care and Use Committee guidelines (IACUC). Ubc-Cre^ERT2/+^ mice^51^ (The Jackson Laboratory) and Atg7^flox/flox^ mice^1^ (provided by Dr. M. Komatsu, Tokyo Metropolitan Institute of Medical Science) were cross-bred to generate the Ubc-Cre ^ERT2/+^; Atg7^flox/flox^ mice as previously described ^2^To generate Ubc-Cre^ERT2/+;^ Atg7^flox/flox^; *Ccl2^-/-^, Ccl2^-/-^* ^23^ (The Jackson Laboratory) were cross-bred with our previously created Ubc-Cre ^ERT2/+^; Atg7^flox/flox^ mice. To generate Ubc-Cre ^ERT2/+^; Atg7^flox/flox^; *Cxcl10^-/-^, CXCL10^-/-^* ^27^ (The Jackson Laboratory) were cross-bred with our previously created Ubc-Cre ^ERT2/+^; Atg7^flox/flox^mice. To Ubc-Cre ^ERT2/+^; Atg7^flox/flox^;L*ep^ob^*/*Lep^ob^,* L*ep^ob^*/*Lep^ob^* ^52^ (The Jackson Laboratory) were cross-bred with our previously created Ubc-Cre ^ERT2/+^; Atg7^flox/flox^mice.

### Tamoxifen Preparation and Administration

TAM (T5648, Sigma) was suspended at a concentration of 20 mg/ml, in a mixture of 98% sunflower seed oil and 2% ethanol. For TAM delivery, 200 μl per 20 g of body weight (20mg/kg) were injected intraperitoneally into 8 to 10 weeks old mice. Mice were treated once per day for 4 days to delete floxed gene systematically^2^. Ubc-Cre^ERT2/+;^ Atg7^flox/flox^; L*ep^ob^*/*Lep^ob^* were treated twice per week for 2 weeks.

### Survival

For mouse Kaplan-Meyer survival curve, mice were monitored daily until they reached the endpoint. The criteria for euthanization were a body condition score of 2, body weight loss of >15%, or natural death.

### Fasting

Fasting was conducted as previous described^2^.

### Metabolic cages

Two indirect calorimetry systems were used, a 12 cage CLAMS apparatus (Columbus Instruments) and 16 cage Promethion Core Mouse Metabolic System (Sable System International). Mice were maintained on a standard chow diet and single housed for 48–72 h prior to experiment start.

During the experiment, mice were single housed under a 12-hour light-dark cycle at 21C and 55 % humidity for 7 days. The first 24 hours of data collection was removed from analysis due to acclimation period. Oxygen consumption, CO_2_ emission, food consumption, movement, running wheel, and energy expenditure were measured every 15 minutes in the CLAMS and 3 minutes in the Promethion.

Locomotor activity, both horizontal and vertical, was determined by a X, Y, and Z infrared light beam system. Stationary locomotor activity was defined as continues infrared light beam breaks of one single light beam and ambulatory movement as continues breaks of two or more different light beams.

Raw data files were collected and processed by the Promethion software package MacroInterpreter 3, which produced standardized output formats for the metabolic variables of interest at each cage. The processed data generated by MacroInterpreter 3 was then analyzed by the CalR: A Web-based Analysis Tool for Indirect Calorimetry Experiments (https://calrapp.org) as described previously^53^.

### Body Composition

Body composition analysis (fat and lean mass) was assessed by the EchoMRI^TM^−100H. Unanesthetized mice were placed in a restraint tube that was inserted into the analyzer for approximately 2 min. The mouse was then returned to its home cage.

### GDF15 ELISA

GDF15 concentration in the serum was determined using a Mouse & Rat GDF-15 ELISA Kit Quantikine ELISA Kit (R&D Systems; MGD150) according to the manufacturer’s instructions.

### Cytokine and chemokine assay

Levels of the secreted cytokines and chemokines were determined using the Procarta Plex® 36-plex immunoassay (Thermo Fischer Scientific; Cat No: EPX360-26092-901) for mouse serum and liver tissue. Data were collected using a Luminex-200 system and validated using the xPONENT software package. Aliquots of serum and tissue in duplicate were assayed for the secreted molecules as per manufacturer’s instructions using Luminex 200 System and analyzed by ProcartaPlex Analyst 1.0 (Luminex Corporation).

### CCL2 ELISA

CCL2 concentration in the liver tissue supernatants was determined using a Mouse CCL2/JE/MCP-1 Quantikine ELISA Kit (R&D Systems; MJE00B) according to the manufacturer’s instructions.

### Production of anti-mCCL2 (C1142)

The complete C1142 mAb (CNTO 888 mouse surrogate) sequence was a kind gift from Janssen Research and Development, LLC. Briefly, the DNA sequences encoding the IgG2a/kappa heavy and light chains of C1142, as well as an irrelevant isotype control mAb were were synthesized by a commercial vendor (GeneArt, Invitrogen), with codon optimization for efficient expression in CHO cells.The ORFs were then sub-cloned separately into customized pTT-based heavy and light chain episomal expression vectors under the control of cytomegalovirus (CMV) promoters. Heavy and light chain vectors were co-transfected into ExpiCHO-S cells (Cat. A29133; Gibco) according to the manufacturer’s instructions and expression allowed to proceed for 5 days. Secreted monoclonal antibodies were purified from clarified expression media using protein A affinity chromatography with MabSelect beads (Cat. GE17-5199-01; Merck), followed by extensive dialysis against phosphate-buffered saline (PBS) using Slide-A-Lyzer G2 dialysis cassettes (Cat. 87731; Life Technologies).

### Serum biochemistry analysis

Blood serum samples were analyzed by the Element DC5X^TM^ Veterinary Chemistry Analyzer (Hesk) performed at Rutgers In Vivo Research Services (IVRS) core facility.

### Bulk RNA-seq analysis

At 8 weeks post deletion, liver tissue from *Atg7^+/+^*, *Atg7^Δ/Δ^*, *Ccl2^−/−^,* and *Ccl2^−/−^;Atg7^Δ/Δ^* were dissected and flash frozen in liquid nitrogen. FastQC v0.11.9 (https://www.bioinformatics.babraham.ac.uk/projects/fastqc/) was used to assess sequencing quality. Reads were first mapped to the mouse genome using HiSat2 v2.2.1^54^. The genomic index along with the list of splice sites and exons were created by HiSat2 using the genome assembly mm10 from ENSEMBL together with the comprehensive gene annotation from mm10 vM23 from Gencode^55^. Gene level counts were computed using Rsubread v2.8.2^56^ (options isPairedEnd = TRUE, requireBothEndsMapped = TRUE, minOverlap = 80, countChimericFragments = FALSE).

The liver tissue was analyzed separately, and genes were filtered out from further analysis if the mean read count across all samples in the tissue was less than 50. This resulted in 10,563, 10,285, and 21,823 genes that went into further analysis of the brown adipose tissue, GNP, and liver data, respectively. DESeq2 v1.34.0^57^ was used to perform differential gene expression analysis. Differentially expressed genes were used for further analysis and visualization. Gene expression heatmaps were generated with pheatmap v1.0.12 (https://cran.r-project.org/web/packages/pheatmap/index.html) using values that were z-score normalized for each gene across all samples within each tissue. Volcano plots were generated with EnhancedVolcano v1.12.0 (https://github.com/kevinblighe/EnhancedVolcano). All analysis starting from count table generation was conducted in the R statistical environment v4.1.3.

### snRNA-seq analysis

At 8 weeks post deletion, hypothalamus tissue from *Atg7^+/+^*, *Atg7^Δ/Δ^*, *Ccl2^−/−^,* and *Ccl2^−/−^;Atg7^Δ/Δ^* were dissected and flash frozen in liquid nitrogen. Downstream analysis was carried out using the scanpy package v1.9.3^58^. Initial quality control steps and normalization were carried out separately for each of the four samples. Cells were filtered out if they had high relative mitochondrial UMI counts (>4-10% for *Atg7^+/+^*, *Atg7^Δ/Δ^*, *Ccl2^−/−^,* and *Ccl2^−/−^;Atg7^Δ/Δ^*) and high total counts (>15,000-20,000 for *Atg7^+/+^*, *Atg7^Δ/Δ^*, *Ccl2^−/−^,* and *Ccl2^−/−^;Atg7^Δ/Δ^*), which resulted in the removal of 150-300 cells for *Atg7^+/+^*, *Atg7^Δ/Δ^*, *Ccl2^−/−^,* and *Ccl2^−/−^;Atg7^Δ/Δ^*. Cells with the potential of being doublets (score >0.2 as detected by Scrublet, 200-250 cells in each sample, respectively) were also removed. Genes were filtered out from the subsequent analysis if they were present in <1% of cells in the sample. Gene expression counts were then normalized with analytical Pearson residual normalization from scanpy, using a theta value of 10 for all four of the samples. After normalization, the four samples were concatenated. Non-protein-coding genes (2,447 genes, 1.6% of total UMI counts) were also filtered out on the basis of the CellRanger mm10 GTF file vM23. This resulted in a dataset of 20,297 cells and1,600 genes.

PCA was run with 100 components, a kNN graph was built using 30 neighbors, 70 PCs and cosine metric, and Leiden clustering was carried out with a resolution of 2.1, resulting in 52 clusters. Known marker genes from HypoMap^58^ were used to annotate the Leiden clusters using the score genes function in scanpy and by exploring differentially expressed genes in each cluster as compared with all cells outside the cluster, obtained using a custom script. For differential expression analysis, log2 fold change (log2FC) of expression was calculated as the ratio of pseudobulk raw UMI counts summed over cells within and outside the cluster (then normalized by total amount of UMI counts inside and outside the cluster), p-values were calculated using Mann-Whitney U test applied to Pearson residual normalized expression values in single cells within and outside the cluster, and Bonferroni correction for multiple hypothesis testing applied to all genes with abs(log2FC) > 0.5.

### Tolerance Test

LL-lactate tolerance tests were performed after 6 h of fasting. Mice were injected intraperitoneally with L-lactate (2 g/kg BW). Blood glucose levels (Accu-Chek Performa glucometer) were determined from the tail vein at 0, 15, 30, 45, 60, and 120 min after injection (Accu-Chek Performa glucometer).

### Histologic and immunohistochemical analysis

Mouse tissues were collected and fixed in 10% formalin solution (Formaldehyde Fresh, Fisher Scientific, SF94-4). Tissues were fixed overnight and then transferred to 70% ethanol for paraffin-embedded sections. The slides were deparaffinized, rehydrated and hematoxylin–eosin staining was performed.

### Metabolite analysis by LC–MS

Metabolites were extracted as described previously^59^. Briefly, metabolites were extracted from serum using the extraction buffer containing methanol: acetonitrile: H_2_O (40:40:20). The final extract was stored at −80 °C until analysis by LC–MS. The LC-MS metabolomic analysis was performed at the Metabolomics Shared Resource of Rutgers Cancer Institute on a Q Exactive PLUS hybrid quadrupole-orbitrap mass spectrometer coupled to a Vanquish Horizon UHPLC system (Thermo Fisher Scientific, Waltham, MA) with an XBridge BEH Amide column (150 mm × 2.1 mm, 2.5 μm particle size, Waters, Milford, MA). The HILIC separation used a gradient of solvent A (95%:5% H_2_O:acetonitrile with 20 mM acetic acid, 40 mM ammonium hydroxide, pH 9.4) and solvent B (20%:80% H_2_O:acetonitrile with 20 mM acetic acid, 40 mM ammonium hydroxide, pH 9.4). The gradient was 0 min, 100% B; 3 min, 100% B; 3.2 min, 90% B; 6.2 min, 90% B; 6.5 min, 80% B; 10.5 min, 80% B; 10.7 min, 70% B; 13.5 min, 70% B; 13.7 min, 45% B; 16 min, 45% B; 16.5 min, 100% B; and 22 min, 100% B^60^. The flow rate was 300 μL/min. The column temperature was set to 25 °C. The autosampler temperature was set to 4 °C, and the injection volume was 5 μL. MS scans were obtained in negative ionization mode with a resolution of 70,000 at m/z 200, in addition to an automatic gain control target of 3 x 10^6^ and m/z scan range of 72 to 1000. Metabolite data was obtained using the MAVEN software package^61^ (mass accuracy window: 5 ppm).

### Labelled Lactate infusion

For intra-jugular vein catheterization, the procedure was performed as described previously^59^. Briefly, venous catheters were surgically implanted into the jugular veins of *Atg7^Δ/Δ^*, *Atg7^+/+^, Ccl2*^-/-^, C*cl2*^-/-^; *Atg7*^Δ/Δ^ mice at 5 weeks post TAM injection. On the day of infusion, mice were fasted for 6 hours. Mice were infused with 13C-Lactate (CLM-1579-PK) dissolved in sterile saline at a rate of 0.1 μL/g/min for 2.5 hours. Mice were sacrificed after infusion for serum analysis by LC-MS.

### Real-time PCR

Total RNA was isolated from hypothalami by Qiagen RNA micro kit (Qiagen). cDNA was then reverse transcribed from the total RNA by MultiScribe RT kit (Thermo Scientific). Real-time PCR were performed on Applied Biosystems StepOne Plus machine using SYBR green master mix (Thermo Scientific). Results were calculated using ΔΔCt method and then normalized to actin.

### Statistical analysis

Statistical analysis was performed with GraphPad Prism (V.6). A Student’s t-test or a one-way analysis of variance (ANOVA) was used for comparison between the groups. A two-way ANOVA was used for repeated measures for comparisons between the groups. A post-hoc comparison using Tukey HSD was applied according to the two-way ANOVA results. Statistical significance was set at p<0.05.

