## Supplementary figures and images for "Autophagy Suppresses CCL2 to Preserve Appetite and Prevent Lethal Cachexia"

### Supplemental Figure 1

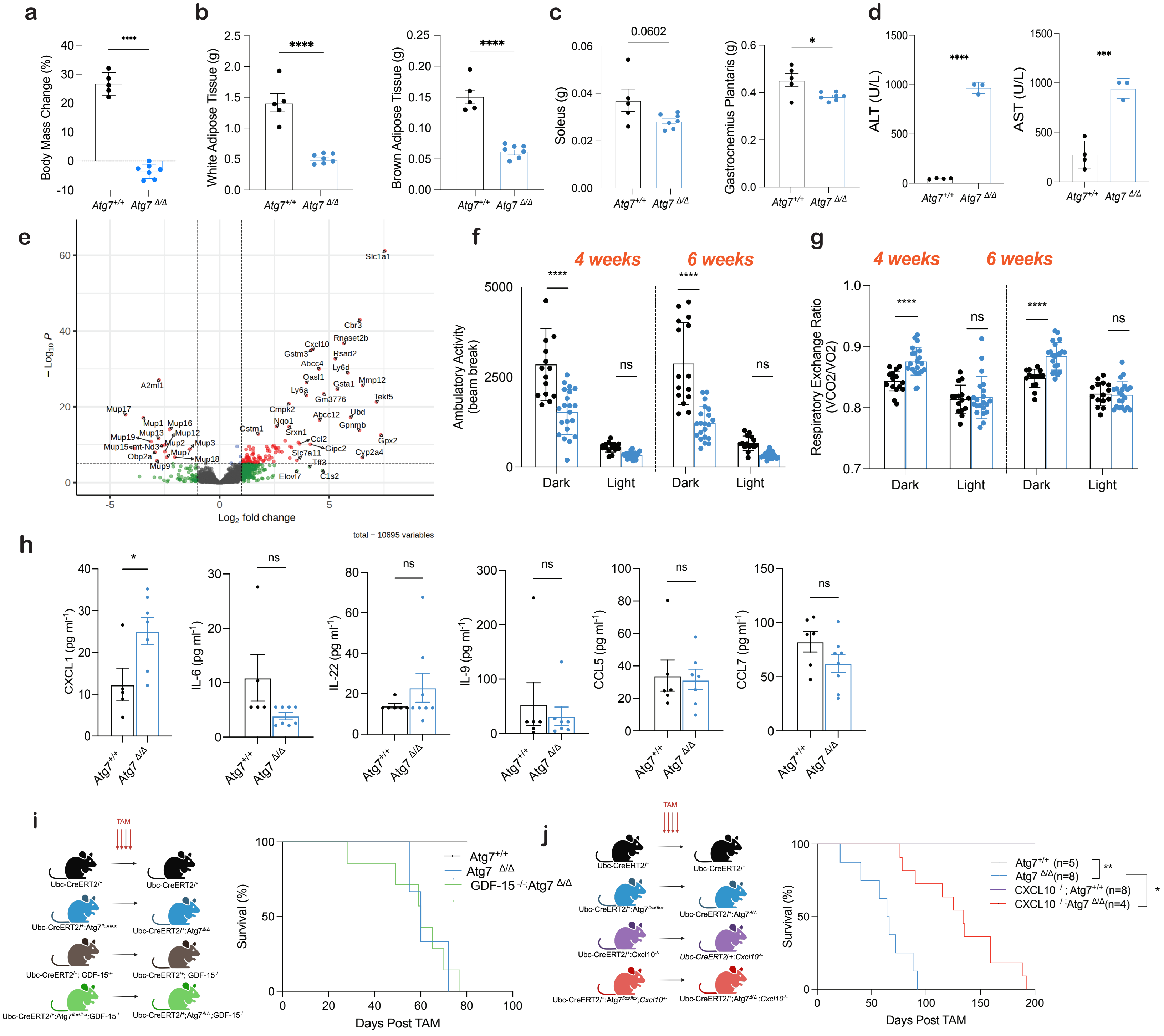

### Supplemental Figure 2

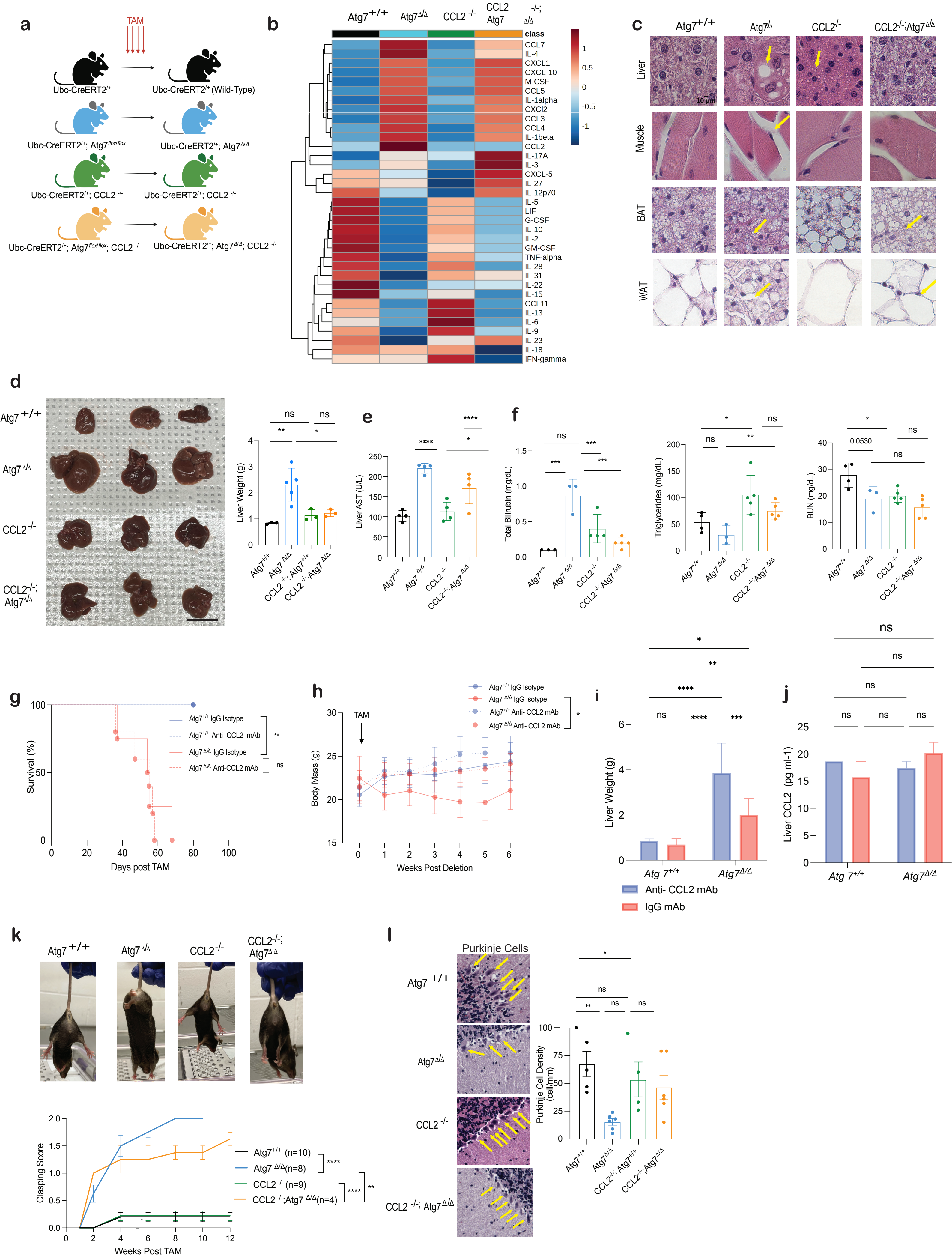

### Supplemental Figure 3

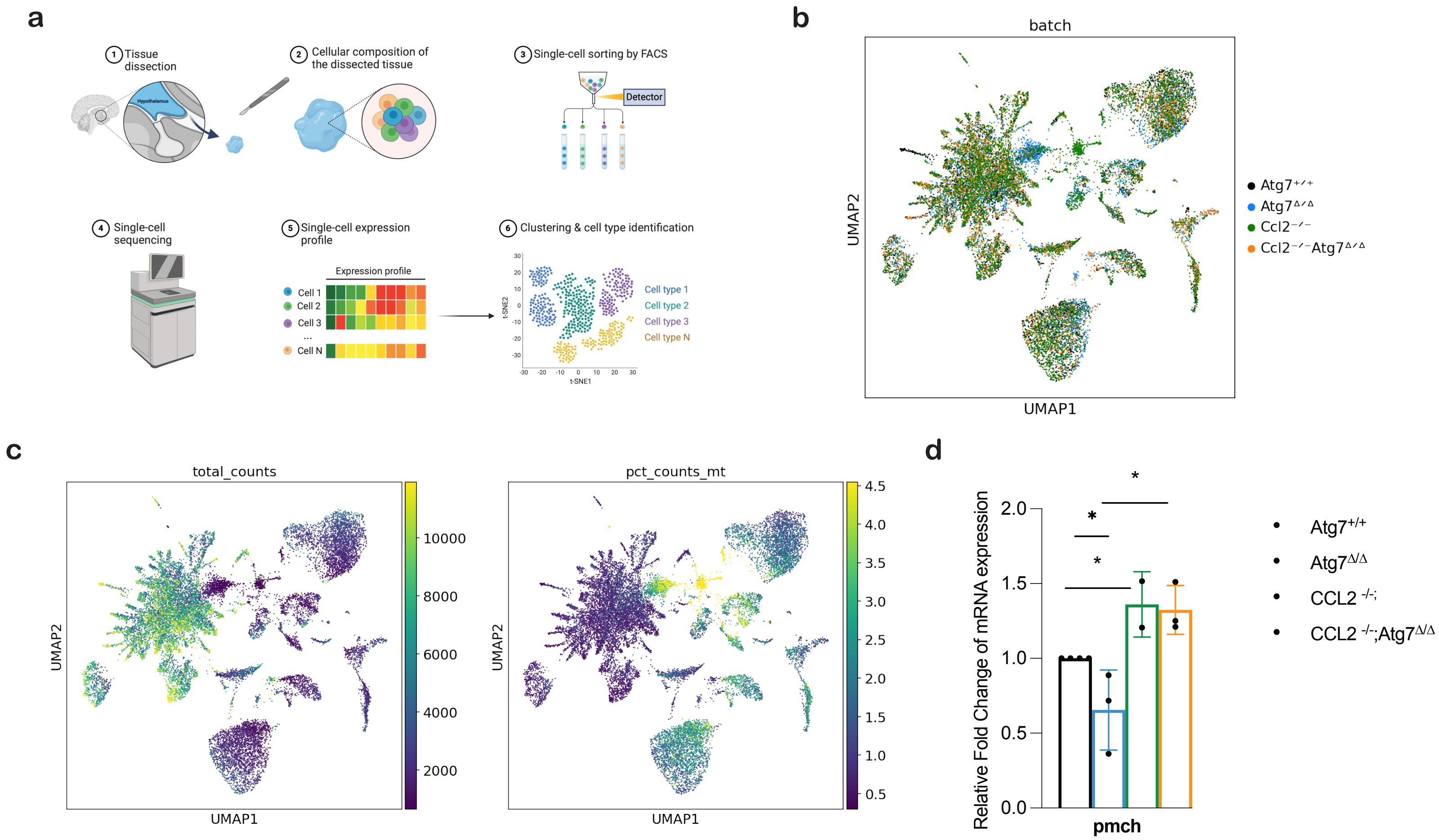
