## Supplementary material for "Autophagy Suppresses CCL2 to Preserve Appetite and Prevent Lethal Cachexia": Legends

**Supplemental Figure 1: Impairment due to loss of *Atg7* causes weight loss and systemic inflammation**

**a**, Percent change from initial body mass of *Atg7^+/+^* mice (*n* = 5) and *Atg7^Δ/Δ^* mice (*n* = 7) at 10 weeks post TAM. **b,c** Mouse tissue weights post TAM injection in *Atg7^+/+^* mice (*n* = 5) and *Atg7^Δ/Δ^* mice (*n* = 7). **d,** Comprehensive Metabolic Panel of ALT and AST levels in serum from *Atg7^+/+^*mice (*n* = 5) and *Atg7^Δ/Δ^* mice (*n* = 3). **e,** Volcano plot depicting differentially expressed genes *Atg7^+/+^* and *Atg7^Δ/Δ^* livers. Y-axis denotes − log10 P values while X-axis shows log2 fold change values. **f-g,** Values are overall hourly means at 4- and 6- weeks post TAM. **f,** ambulatory acivity **g,** RER. **h,** Serum cytokine and chemokine profiling (*n* = 3–11/group) of *Atg7*^+/+^ and *Atg7*^Δ/Δ^ mice. **i,** Experimental design for generation of *Atg7^+/+^*, *Atg7^Δ/Δ^* mice, *Gdf15^-/-^* mice, and *Gdf15^-/-^;Atg7^Δ/Δ^* mice. Ubc-Cre*^ERT2/+^*, Ubc-Cre*^ERT2/+^; Atg7^flox^*^/^*^flox^* mice, Ubc-Cre*^ERT2/+^; Gdf15^-/-^*, Ubc-Cre*^ERT2/+^; Gdf15^-/-^; Atg7^flox^*^/^*^flox^* mice were treated with TAM at 8–10 weeks of age. Kaplan-Meier survival curve of *Atg7^+/+^* , *Atg7^Δ/Δ^* , *Gdf15^-/-^*, and *Gdf15^-/-^* ;*Atg7^Δ/Δ^* mice. **j,** Experimental design for generation of *Atg7^+/+^*, *Atg7^Δ/Δ^* mice, *Cxcl10^-/-^* mice, and *Cxcl10^−/−^;Atg7^Δ/Δ^* mice. Ubc-Cre*^ERT2/+^*, Ubc-Cre*^ERT2/+^; Atg7^flox^*^/^*^flox^* mice, Ubc-Cre*^ERT2/+^; CXCL10^-/-^*, Ubc-Cre*^ERT2/+^; CXCL10^-/-^; Atg7^flox^*^/^*^flox^* mice were treated with TAM at 8–10 weeks of age. Kaplan-Meier survival curve of *Atg7^+/+^*, *Atg7^Δ/Δ^* , *Cxcl10^-/-^*, and *Cxcl10^−/−^;Atg7^Δ/Δ^* mice.

**Supplemental Figure 2: Loss of CCL2 rescues tissue damage, not treatment with C1142 mAb**

**a,** Experimental design for generation of *Atg7^+/+^*, *Atg7^Δ/Δ^* mice, *Ccl2^-/-^* mice, and *Ccl2^−/−^;Atg7^Δ/Δ^* mice. Ubc-Cre*^ERT2/+^*, Ubc-Cre*^ERT2/+^; Atg7^flox^*^/^*^flox^* mice, Ubc-Cre*^ERT2/+^; Ccl2^-/-^*, Ubc-Cre*^ERT2/+^; Ccl2^-/-^; Atg7^flox^*^/^*^flox^* mice were treated with TAM at 8–10 weeks of age. **b,** Heat map of serum cytokine and chemokine profiling in *Atg7^+/+^*, *Atg7^Δ/Δ^* mice, *Ccl2^-/-^* mice, and *Ccl2^−/−^;Atg7^Δ/Δ^* mice at 8 weeks post deletion. **c,** Representative histology of liver, muscle, white adipose tissue (WAT), and brown adipose tissue (BAT) by hematoxylin and eosin stain (H&E) from *Atg7^Δ/Δ^* mice, *Ccl2^-/-^* mice, and *Ccl2^−/−^;Atg7^Δ/Δ^* mice at the 8-wk time point. Yellow arrows indicate the damage site for these tissues. **d,** Representative image of livers from *Atg7^+/+^*, *Atg7^Δ/Δ^*, *Ccl2^-/-^*, and *Ccl2^−/−^;Atg7^Δ/Δ^* mice at the 8-wk time point. Measurement of liver weights from *Atg7^+/+^*, *Atg7^Δ/Δ^*, *Ccl2^-/-^*, and *Ccl2^−/−^;Atg7^Δ/Δ^* mice at the 8-wk time point (*n*=3-5/group). **e-f,** Comprehensive Metabolic Panel serum levels of from *Atg7^+/+^*, *Atg7^Δ/Δ^*, *Ccl2^-/-^*, and *Ccl2^−/−^;Atg7^Δ/Δ^* mice at the 8-wk time point (*n*=3-5/group). **e,** liver AST levels **f,** total bilirubin**,** triglycerides**, and** BUN levels. **g,** Kaplan-Meier survival curve of *Atg^+/+^*  and  *Atg7^Δ/Δ^* mice on Anti-CCL2 mAb or IgG mAb. **h,** Measurement of body weight over 6 weeks of treatment. **i,** Measurement of liver weights at 6 weeks of treatment. **j,** Liver CCL2 ELISA. **k,** Representative images of hindlimb clasping from *Atg7^+/+^*, *Atg7^Δ/Δ^*, *Ccl2^-/-^*, and *Ccl2^−/−^;Atg7^Δ/Δ^*  mice at the 8-wk time point and clasping score. **l,** Representative histology of cerebellum by hematoxylin and eosin stain (H&E).Yellow arrows indicate the Purkinje cells in these tissues. The density area of Purkinje cells were quantified in panel (*n*=3 per group).

**Supplemental Figure 3: Reduced mRNA expression of PMCH during loss of autophagy.**

**a,** Schematic of the scRNA-seq workflow. **b,** UMAP overlay of *Atg7^Δ/Δ^* mice, *Ccl2^-/-^*, and *Ccl2^−/−^;Atg7^Δ/Δ^* samples at the 8-wk time point. **c,** UMAP of total snRNA-seq read counts and percentage of mitochondrial gene expression counts. **d,** Relative fold change mRNA expression levels of pmch in hypothalamus through qRT-PCR from *Atg7^Δ/Δ^*, *Ccl2^-/-^*, and *Ccl2^−/−^;Atg7^Δ/Δ^* mice at the 8-wk time point. (*n*=3-4/group) .
